# MoleMap: fast alignment-free molecule mapping for long-read and linked-read sequencing data

**DOI:** 10.1101/2022.06.20.496811

**Authors:** Richard Lüpken, Thomas Krannich, Markus Schuelke, Birte Kehr

## Abstract

**Motivation:** The bottleneck for genome analysis will soon shift from sequencing cost to computationally expensive read alignment. When only portions of the genome are of interest, read alignment on whole-genome sequencing (WGS) data can be replaced by a fast mapping step.

**Results:** We propose an approach that can map both long reads or linked-read molecules, as a pre-processing step for downstream targeted analyses. Our “molecule mapping” approach, implemented in the tool *MoleMap*, uses a minimized open-addressing *k*-mer index of the reference genome and a fast *k*-mer clustering procedure. We demonstrate that MoleMap’s accuracy is competitive with standard alignment and mapping tools and its running time outperforms other mapping tools on 32 threads by a factor of 3 to 8 and read alignment tools by a factor of 10 to 60. Its low memory footprint allows us to analyze whole genomes on a standard laptop computer. As proofs of concept, we use MoleMap to filter reads for local assembly of a known variant region that involves non-reference sequence and showcase its use in diagnosing a patient with a rare disease. Our work contributes to more scalable genome analysis and promotes WGS for targeted analyses.

**Availability and Implementation:** Source code of MoleMap is available at https://github.com/kehrlab/molemap.

## Introduction

Genomic analyses frequently focus on selected regions of the genome, for example, few target loci [Jang et al., 2018], small gene panels [Viswanathan et al., 2018, Greer et al., 2017] or exonic regions only [Akbari et al., 2021]. As costs for sequencing drop and the 100$ genome comes into reach [Pennisi, 2022], WGS may soon replace panel and whole-exome sequencing for a range of applications. Full WGS provides the flexibility to perform a targeted analysis while retaining the option to incrementally expand the analysis from few and shorter to more and larger target regions, as the data set itself is not limited to a subset of the studied genome. However, whole-genome data sets are substantially larger than data from targeted sequencing approaches. Handling these large amounts of data can quickly exceed available computational power and thus can hinder the decision to generate whole-genome data. For fully transitioning from panel and whole-exome sequencing to WGS, data analysis needs to become computationally more efficient.

In addition to transitioning towards whole-genome data, the sequencing field is currently moving rapidly from short to long reads [Kovaka et al., 2023]. Over the last decade, the error rates of long reads have improved substantially to a level that is now competitive with short reads. In particular, PacBio HiFi reads [Wenger et al., 2019] offer long read lengths combined with error rates well below 1%. We can now fully leverage the long-range information provided in long-read WGS data, on the one hand to resolve complex genomic regions [Nurk et al., 2022], and on the other hand also to tackle the computational bottleneck when analyzing whole-genome data as we show below. Similar opportunities for leveraging long-range information exist for data from linked-read technologies [Zheng et al., 2016, Wang et al., 2019, Chen et al., 2020]. Essentially, linked reads are short reads labeled with barcodes providing long-range information over distances like long reads, while retaining the relatively low cost of short reads. Reads with the same barcode originate from a small set of large DNA molecules. This equips the data with an additional level of structure, which we can harness for reducing computational requirements.

Read-to-reference alignment typically constitutes the largest portion of the computational, and consequently also of the monetary cost of genome analysis. Therefore, replacing the expensive alignment with a fast alignment-free mapping approach can substantially speed up the analysis and may eventually be necessary to establish WGS for a wider range of analyses. For long reads, minimap2 [Li, 2018] has become the alignment software of choice. Recent improvements to the computational efficiency of minimap2 were made by BLEND [Firtina et al., 2023], which directly modifies the minimap2 software and speeds up the alignment of HiFi reads severalfold without affecting accuracy. In addition, minimap2 itself is capable to map long reads omitting base-pair precise alignment for reducing the computational load. Similarly, MashMap [Jain et al., 2018] and mapquick [Ekim et al., 2023] provide a mapping functionality. The tool mapquick is orders of magnitude faster than minimap2 but designed for low-divergence mapping without full read alignment.

Aiming for even greater computational efficiency, we here suggest an alignment-free read filter that omits full whole-genome read alignment, thereby enabling subsequent, detailed analyses on genomic regions of interest. We introduce a mapping approach that can equally assign linked-read barcodes or long reads to genomic intervals. We refer to our approach as *molecule mapping*, where a molecule can either be a single long read or a set of linked reads labeled with the same barcode. The implementation in our software *MoleMap* allows us to extract reads from regions of interest in a fraction of the time needed for full read alignment. Mapping can be followed by read alignment or local assembly, limited to genomic regions of interest and thus having negligible running time. In our evaluations, MoleMap achieves severalfold speedups over popular mapping and alignment tools with mostly minor trade-offs in accuracy. As opposed to other work focusing on alignment speed [Miller et al., 2015], we achieve this speed-up on commodity hardware. Requiring less than 8 GB of RAM on human genome data MoleMap can even be run on a standard laptop. Finally, we demonstrate how targeted analyses can benefit from the tremendous savings in computational time and offer guidance on the expected speed-up compared to full read alignment depending on the size of genomic regions of interest.

## Materials and methods

MoleMap takes as input the reference genome, and a long-read FASTQ file or a barcode-sorted pair of linked-read FASTQ files. MoleMap first builds a *k*-mer index of the reference genome, which is analogous to the index files used by popular read aligners. In the main mapping step (Figure 1), MoleMap predicts the molecules’ intervals of origin in the reference genome. Mapping locations and scores are written to a file, either for all reads or only for reads falling within user-specified genomic regions. If mappings of all reads are written to the output file, subsequent sorting and indexing enables efficient and incremental retrieval of reads originating from regions of interest.

**Fig. 1.**
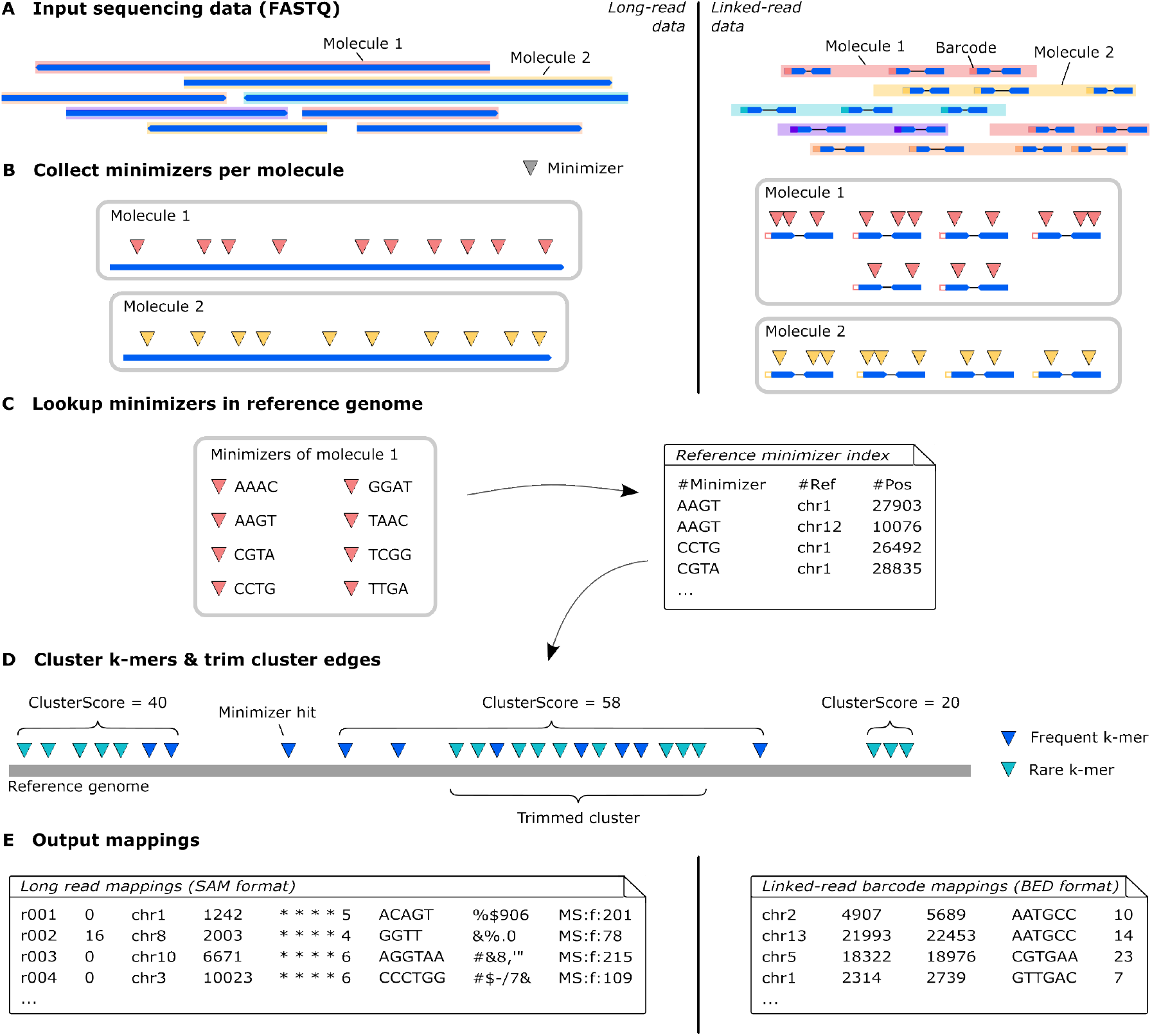
Overview of the molecule mapping workflow implemented in MoleMap. The left side shows the workflow for long reads while the rigth side shows the workflow for linked-read. Steps that are centered in the figure (C and D) are common to both data types differing only in default parameters values.

### Constructing the MoleMap index

Key to the comparably low memory footprint of MoleMap is its index data structure. MoleMap builds a minimized open-addressing *k*-mer index of the reference genome. An open-addressing index allows for ambiguity when hashing the *k*-mers [Weese, 2013] and thereby massively reduces the required memory for large values of *k* (e.g. *k* = 17). While the size of a standard *k*-mer index scales with the number of possible *k*-mers, the size of an open-addressing index scales only with the length of the indexed sequence. The size advantage comes at the cost of slightly longer look-up times for resolving hash collisions. MoleMap resolves a collision by performing predefined bit-operations on the hash value until a valid index position is found (Supplementary Figure 1, Supplementary Section 1.1).

In addition to using open-addressing, MoleMap uses a minimizing strategy for constructing the index. We chose a window-based minimizing technique similar to minimap [Li, 2016] where the *k*-mer with the lowest hash value within a fixed size window of *w* consecutive *k*-mers is defined as the window’s minimizer. Only the *k*-mers that are minimizers of some window of size *w* are stored in the index (default: *k* = 17, *w* = 9 for noisy long reads; *k* = 19, *w* = 19 for HiFi long reads; and *k* = 31, *w* = 31 for linked reads). To avoid a bias towards minimizer sequences such as poly-A and reduce collisions during index look-ups, MoleMap aims for a random *k*-mer order by performing a bitwise *xor* operation between the bit-vector representation of the *k*-mer and a consistently reused pseudo-random bit-vector of length 2*k*.

For faster read analysis in the molecule mapping step, MoleMap uses an index design that allows to look up the frequency of a minimizer without accessing all its positions while maintaining a minimal memory footprint (Supplementary section 1.2).

### Mapping long reads with MoleMap

We first describe the mapping of long reads before introducing differences for linked reads. Wherever we refer to a long read as a molecule in this subsection, the described approach applies equally to linked-read data.

Similarly to read aligners that align reads independently of each other, MoleMap maps each molecule independently of other molecules (Figure 1A). For each molecule, MoleMap determines all its minimizers (Figure 1B) and looks them up in the *k*-mer reference index (Figure 1C). If the abundance *A*_*m*_ in the reference genome of a minimizer *m* is sufficiently small (default ≤ 20), MoleMap extracts its reference positions *P*_*m*_ = {*p*_*m*_} from the index and adds them to a list of *minimizer hits M*. In addition, MoleMap records for all minimizers *m* in the molecule their abundance *A*_*m*_ in the reference genome and determines the number *B*_*m*_ of consecutive windows in the read sequence for which *m* remains the minimizer. Only for long reads, MoleMap further stores the read position *o*_*m*_ of each minimizer *m*. Once a complete list of minimizer hits *M* is created for a molecule, MoleMap sorts the list by position on the reference genome.

MoleMap clusters minimizer hits in *M* using two criteria and a cluster score (Figure 1D):

- The *maximum-range criterion* requires hits of the same cluster *C* ⊆ *M* to originate from the same reference contig (chromosome) and limits the range a cluster spans in the contig by an upper threshold (default 300,000).
- The *maximum-gap-size criterion* limits the distance between two adjacent hits in a cluster to a maximum size (default 20,000 chosen empirically; see Supplementary Figure 5, Supplementary Section 3.5).
- MoleMap accumulates a *cluster score S*(*C*) from all minimizer hits within a cluster, *p*_*m*_ ∈ *C*,

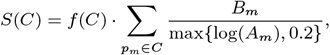

where *f* (*C*) is a scaling factor that differs between long reads and linked reads as defined below. The number *B*_*m*_ of consecutive windows for which a minimizer *m* stays the same, linearly increases the contributed score. Rare minimizers contribute more than frequent minimizers to the score, scaling inverted logarithmically as a function of abundance *A*_*m*_.

For long reads, we define the *scaling factor f* (*C*) to account for the order of minimizer hits in the reference compared to the read, so that *f* (*C*) = 1 if the order is the same in reference and read, and 0 ≤ *f* (*C*) *<* 1 if the order of some minimizers is inconsistent (Supplementary section 1.3). Although using our scaling factor is less accurate than the longest increasing subsequence algorithm [Schensted, 1961] as used in minimap [Li, 2016] or a chaining algorithm [Ohlebusch and Abouelhoda, 2005] as used in minimap2 [Li, 2018], it is fast and effective.

MoleMap identifies initial clusters that conform with the maximum-range and maximum-gap-size criteria and maximizes the cluster score by iterating over the sorted list of minimizer hits. Hits are added to a cluster as long as both criteria are met. If the maximum range criterion is violated, multiple cluster configurations spanning the same genomic region may be possible. In this case, MoleMap chooses the cluster with the highest cluster score. If the maximum gap size criterion is hurt, the current cluster is saved and a new cluster is initialized with the new minimizer hit. We note that this simple clustering procedure is heuristic and does not optimize a global score.

After the initial clustering, MoleMap post-processes every cluster (Figure 1D) with the aim of removing false expansions of mappings that are caused by spurious random minimizer hits. More specifically, MoleMap applies a sliding window running inward from the start and another from the end of the initial cluster. These sliding windows check if a *cluster-score-per-base threshold* (default 20 per 4000 *bp* window for long reads and 100 per 2000 *bp* window for linked reads) is met for the edges of the cluster and, if necessary, trim the set of clustered minimizer hits until this condition is met at both ends. Only clusters that are longer than a certain length (default 100 for long and 10000 for linked reads) are kept after edge trimming. MoleMap does not pre-define a lower bound on the mapping score *S*(*C*), but it can be set by the user. By creating a mapping score histogram, MoleMap provides guidance in choosing a minimum score threshold that is specific to the sample, the parameters and the reference genome (Supplementary Figure 6, Supplementary Section 3.6).

Finally, MoleMap converts all clusters into molecule mappings. For long reads, MoleMap outputs them in the SAM format [Li et al., 2009], leaving the CIGAR string empty. The mapping score is provided in the MS tag.

### Mapping and extracting linked reads with MoleMap

With linked-read technologies, a long DNA molecule is sequenced as a set of short reads that share the same barcode. MoleMap processes linked reads per barcode, i.e., it computes mappings for each set of reads labeled with the same barcode. As some linked-read technologies attach the same barcode to reads from several long DNA molecules, we acknowledge that it would be more accurate to refer to our approach for linked-read data as barcode mapping rather than molecule mapping. By reporting several mappings per barcode, MoleMap is able to handle several long DNA molecules from different regions of the genome simultaneously.

MoleMap’s processing steps for linked- and long-read data are largely identical. The main difference between long and linked reads is the input and output (Figure 1), the values of the default parameters, and the process for retrieving reads from regions of interest. For linked reads, MoleMap requires the input pair of FASTQ files to be sorted by barcode. MoleMap does not map individual reads from linked-read data sets but accumulates the list of minimizer hits *M* over all reads per barcode (Figure 1B). When mapping linked reads, the scaling factor *f* (*C*) accounts for the span of the clusted minimizer hits over the reference (Supplementary Section 1.3). Instead of writing a SAM file, MoleMap writes a BED-formatted output file for linked reads. For each mapping, MoleMap’s BED file stores the reference contig, start and end position, the mapped barcode, and the mapping score. After sorting the BED file by genomic position, extracting all barcodes mapped to a user-defined region is straightforward, e.g., using BEDtools [Quinlan and Hall, 2010]. MoleMap additionally writes a read index file that stores the positions of the barcodes in the input FASTQ files. Using this index, reads from genomic regions of interest can then be efficiently retrieved. For linked reads, MoleMap additionally offers an adaptive score threshold that can be used optionally for filtering mappings (Supplementary Figure 2, Supplementary Section 1.4).

### Implementation

The MoleMap software tool implements four commands: `MoleMap index` creates the *k*-mer index of the reference genome; `MoleMap mapLong` and `MoleMap mapLinked` compute the mapping of molecules to the reference genome using the reference index; and `MoleMap get` can be used to retrieve linked reads labeled with specific barcodes. For linked reads, we additionally prepared a Snakemake workflow [Mölder et al., 2021] that chains the MoleMap commands with standard unix commands, offering users a simple way to retrieve reads from regions of interest (Supplementary Section 2.3).

## Results

Here we assess MoleMap’s computational performance on real data, evaluate its mapping accuracy on simulated data, compare its mappings to full alignment results on real data, and describe two example use cases. Throughout all evaluations, we use T2T-CHM13v2.0 as the reference genome unless otherwise stated and compare molecule mappings from MoleMap to read alignments from short-read, linked-read and long-read aligners. If available, we also ran aligners in a “mapping only” mode. We acknowledge that full read alignments contain more information than molecule mappings. Nevertheless, we argue that comparison with aligners is justified in the context of targeted genome analyses from WGS data, since genome-wide alignment is the task that we suggest replacing using MoleMap.

### MoleMap is substantially faster than read aligners on real data

We evaluate MoleMap’s computational performance on publicly available and in-house patient data sets and compared it to popular mapping and alignment tools (Supplementary Sections 2.2).

Molecule mapping with MoleMap proves to be substantially faster than full read alignment, outperforming the fastest aligner at least 8-fold on long reads (Figure 2A,B) and 11-fold on linked reads (Figure 2C). Notably, MoleMap also outperforms minimap2 and Strobealign [Sahlin, 2022] in their mapping modes by factors of 3 and 4.5 respectively. Visualizing the time savings in a linked-read workflow, which includes pre-processing and sorting steps (Supplementary Section 2.3), demonstrates that this speed-up shifts the bottleneck away from read mapping to other steps (Figure 2D). Overall, MoleMap’s running times demonstrate the enormous potential to save time when molecule mapping is sufficient and genome-wide read alignment is not required.

**Fig. 2.**
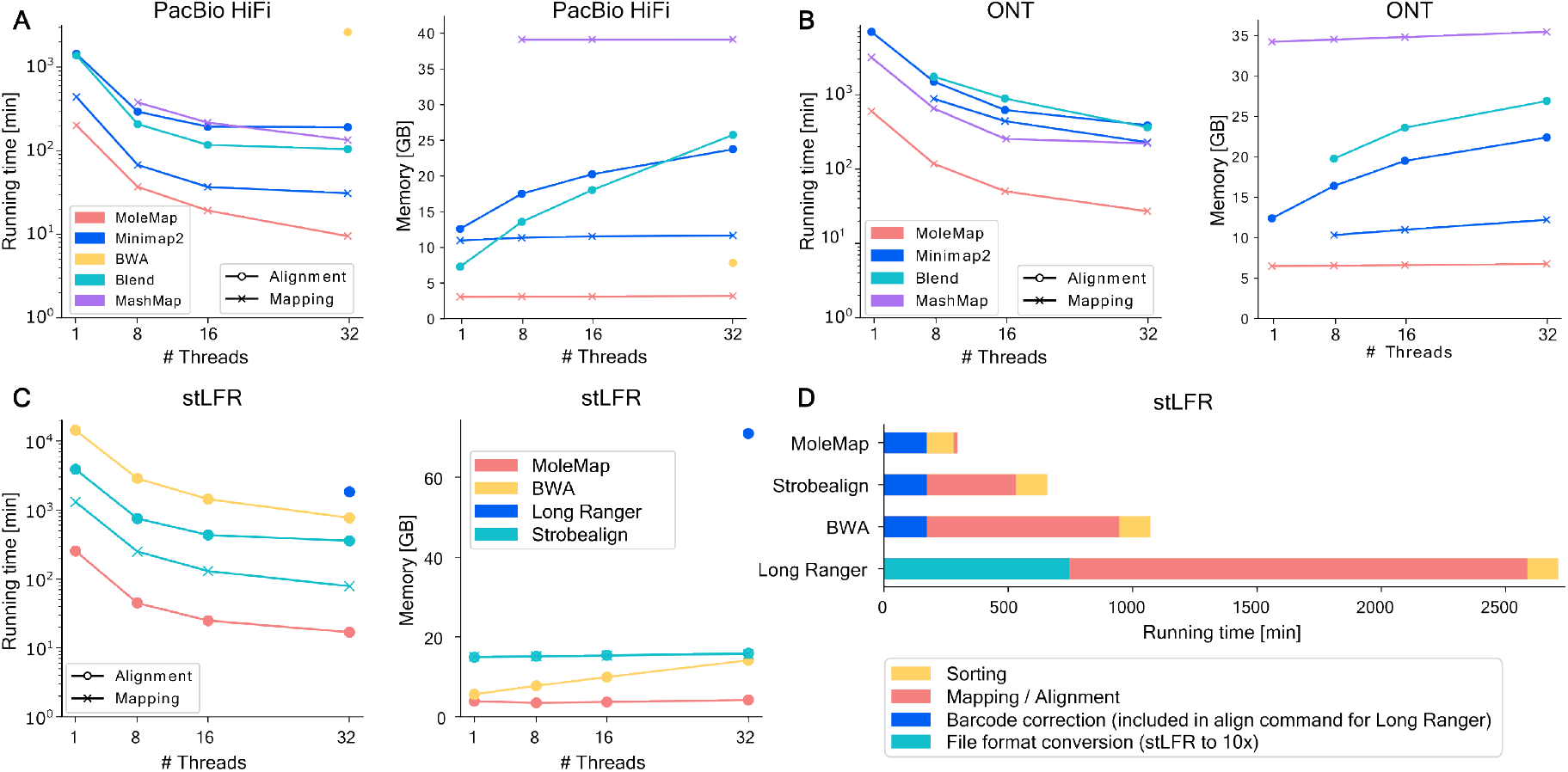
Comparison of running time and memory consumption on real long-read data (A,B) and real linked-read data (C), and workflow running times on 32 threads on the linked-read data (D). Exact values in Supplementary Section 3.2

### MoleMap runs on a laptop due to its low memory footprint

We used the same publicly available and in-house patient data sets to evaluate MoleMap’s memory requirements. MoleMap outperforms the competing tools, requiring less than 7GB of RAM for long reads (Figure 2A,B) and less than 3GB for linked reads (Figure 2C) while scaling well when using more threads. Given the low memory footprint, we tested running MoleMap on a standard laptop with an 8-core CPU and 16 GB of RAM (Supplementary Section 2.2) on public long-read data sets (Supplementary Sections 2.1.1). Building the long-read index took 16:25 min and required 6.78GB of memory. Mapping 4 million PacBio HiFi reads totaling 65.1 Gbp on 7 threads took 52 min and required 5.57GB of memory. Mapping 1 million ONT ultra-long reads totaling 70.4 Gbp on 4 threads took 1:59 hours and required 6.77GB of memory.

### MoleMap achieves high precision and recall on simulated data

Simulated data allows us to directly measure mapping accuracy based on a truth set, i.e., the original genomic positions of simulated reads are known. We simulated various long-read data sets using PBSIM3 [Ono et al., 2022] (Supplementary Section 2.1.3) and linked-read data sets with LRSIM [Luo et al., 2017] (Supplementary Section 2.1.4). Again, we benchmark MoleMap on these data against popular mapping and alignment tools (Supplementary Sections 2.2.2 and 2.2.4). We recorded precision-recall curves (Supplementary Section 3.1) for the simulated long-read data sets by varying mapping score thresholds and for the simulated linked-read data by varying the minimum number of reads per read cluster (Figure 3).

**Fig. 3.**
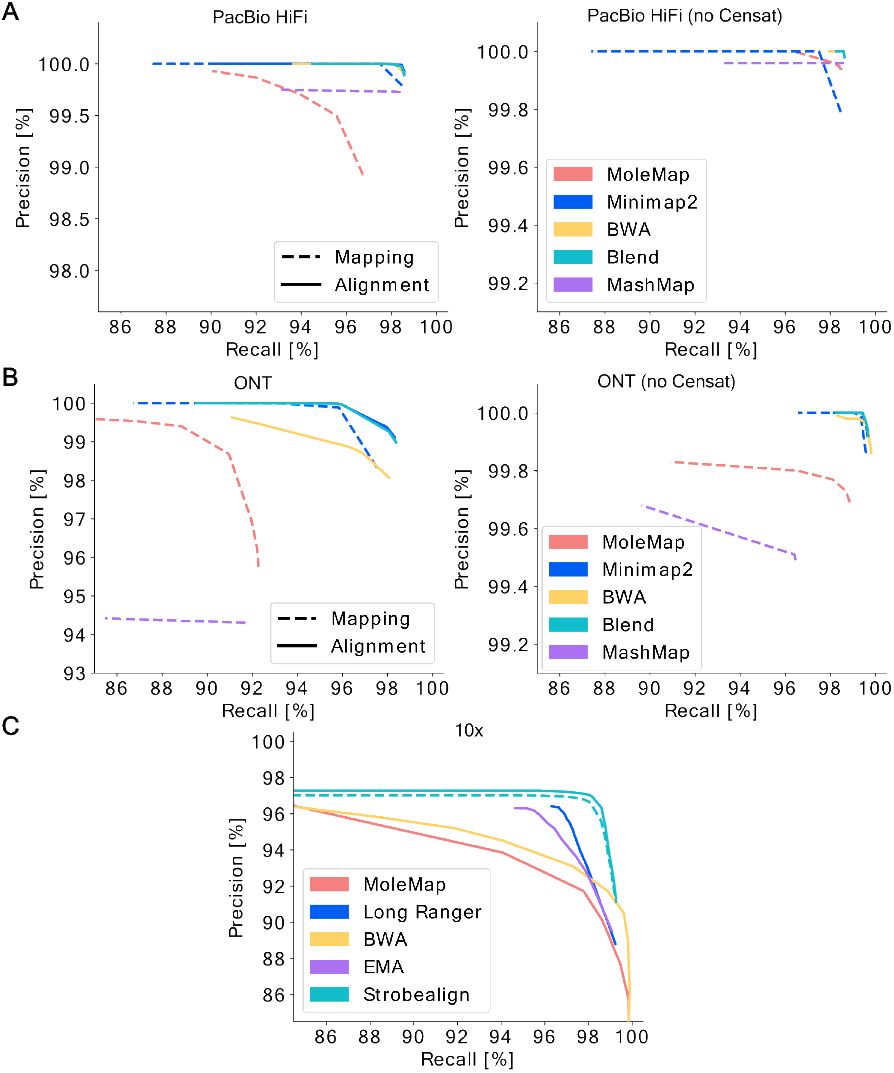
Recall and precision on simulated PacBio HiFi long reads (A), ONT long reads (B) and 10x linked reads (C).

On long reads, MoleMap’s precision is competitive, while slightly lower than that of full read alignment. MoleMap’s recall is several percentage points lower than that of full alignment. We found that most of this is due to centromere and satellite repeat (CenSat) regions specific to the T2T genome. When CenSat regions are excluded from evaluation (Supplementary Section 2.2.5), MoleMap’s recall is only marginally lower than that of the other tools. Both recall and precision of MoleMap are higher on the more accurate PacBio HiFi data (Figure 3A) than on ONT long reads (Figure 3B). Minimap2 and BLEND achieve the best results with almost identical output.

On linked reads, we had to use a simple clustering procedure to infer the truth set of barcode mappings from the data (Supplementary Section 2.1.5). MoleMap achieves recall rates similar to those of the tested short- and linked-read aligners (Figure 3C) and is insensitive to introduced variation (Supplementary Figure 4). While MoleMap’s mapping precision is similar to BWA’s precision, Strobealign and the dedicated linked-read aligners EMA and Long Ranger^1^ outperform MoleMap.

### MoleMap agrees with aligners to a large extent on real data

We also assess the agreement between MoleMap and the competing tools on real data (Supplementary Sections 2.1.1 and 2.1.2).

On PacBio HiFi data, MoleMap and Minimap2 alignments report the same mapping for 97.31% of the reads. For 2.23% of the reads, MoleMap reports another mapping position, and 0.44% of the reads remain unmapped by MoleMap. If we exclude CenSat regions from the assessment, we observe an agreement of 99.68% and a disagreement on the remaining 0.32% of reads. All reads outside of CenSat regions are mapped. The results for minimap2 in mapping mode and BLEND were very similar, whereas the comparison to MashMap shows a lower agreement of just 96.91% and 98.03% without CenSat regions.

Consistent with higher error rates, we observe a slightly lower agreement of 94.52% on the ONT data, and a greater portion of 2.05% unmapped reads by MoleMap and 0.66% unmapped by Minimap2 and BLEND. The remaining 3.36% of reads were mapped to different positions. When excluding CenSat regions, we again observe a higher agreement of 96.47% and only 1.1% of disagreeing mappings, however the number of unmapped reads remains identical for both MoleMap and Minimap2. This indicates that the unmapped reads in this case are caused by the lower base-calling accuracy of the ONT long reads.

On linked-read data, the total number of sequenced molecules is unknown, so we compare the number of mappings reported per tool that agree or disagree with another tool (Figure 4C,D). Between 89.2% and 96.1% of MoleMap mappings were supported by the competing alignments on stLFR data and 90.2% to 91.3% of mappings agreed with competing aligners on the TELL-Seq data. When excluding CenSat regions, MoleMap reaches an agreement between 92.3% and 97.6% on stLFR and 94.1% to 95.1% on TELL-Seq data. We also compare MoleMap to Strobealign in “mapping only” mode. Here we observe a high agreement of 96.1% on the stLFR data, but on the TELL-Seq data Strobealign’s “mapping only” mode proves unreliable reaching an agreement of only 46.0%.

**Fig. 4.**
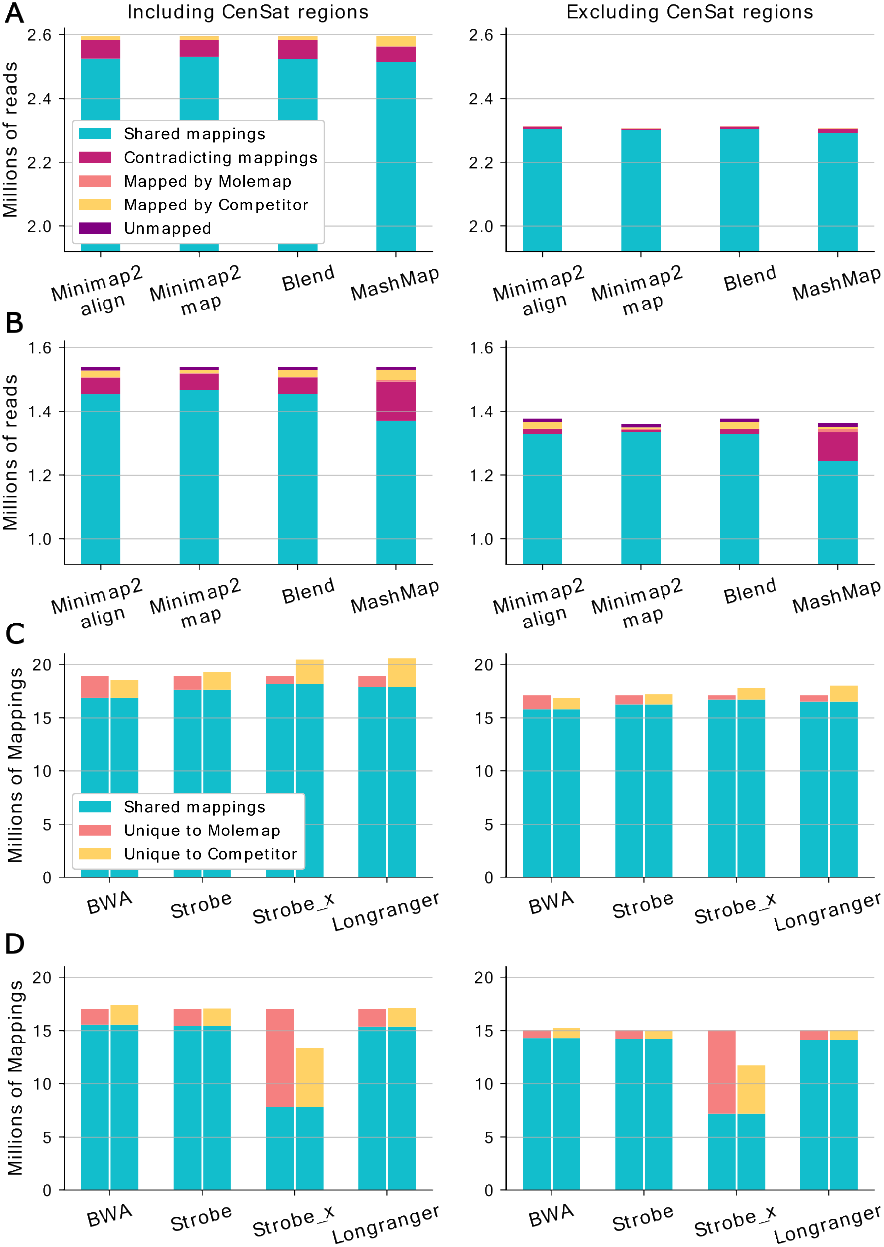
Agreement of MoleMap with competing tools on real PacBio HiFi (A) and ONT (B) long reads and on real stLFR (C) and TELL-Seq (D) linked reads.

### Use case 1: From FASTQ to diagnostic SVs in less than 1 hour

We generated ONT long-read sequencing data for a patient with both intellectual disability and muscular dystrophy carrying two previously identified diagnostic SVs (Supplementary Section 2.1.1). After basecalling, we analyzed the data with a targeted SV calling workflow, which limits genome analysis to regions around a given set of genes (Supplementary Section 2.4.2). In short, the workflow extracts a region around each gene (*±*1000 bp) before using MoleMap for mapping long reads and reporting mappings only in these regions. Subsequently, the remaining reads are aligned with Minimap2 to the reference genome, sorted and indexed with samtools [Danecek et al., 2021], and input to Sniffles [Smolka et al., 2024] for SV detection. For our use case, we compiled a list of 52 genes known to affect the development of intellectual disability and 61 genes associated with muscular dystrophies (Supplementary Section 2.4.1). The workflow detected a total of 162 SVs, including the two previously identified diagnostic deletions with high read support (Supplementary Section 2.4.3). The first deletion, reported by Sniffles at chr12:115,966,387-116,114,816 (148429 bp), overlaps 17 exons of the MED13L, a gene well-known for intellectual disability [Asadollahi et al., 2013, Adegbola et al., 2015]. The second deletion, detected at chrX:32,083,165-32,248,141 (164976 bp), deletes 11 exons of the DMD gene likely causing Duchenne muscular dystrophy [Hoffman et al., 1987]. The running time of the entire targeted variant calling workflow was 34:37 min using 32 threads, of which 31:59 min are spent on molecule mapping. This is substantially faster than the 5:36 h needed by a standard workflow with full read alignment.

### Use case 2: Local assembly of non-reference sequence

We tested whether MoleMap can be applied as the initial step of a local sequence assembly workflow for a given target region of the human genome. We selected the genomic position chr17:17,831,079 on the GRCh38 human reference genome that was previously found to be the breakpoint of a 766 bp non-reference sequence variant associated with a lower risk of myocardial infarction [Kehr et al., 2017]. The Genome-in-a-Bottle consortium previously reported NA12878 to be a carrier of this variant [Zook et al., 2020].

We applied MoleMap to the linked-read data of NA12878 and extracted the set of reads labeled with barcodes that had a mapping overlapping a 50 kbp window around the variant position, i.e., 25 kbp to each side (Supplementary Section 2.5.1). The set of reads was passed to the Minia assembler [Chikhi and Rizk, 2013] of the GATB library [Drezen et al., 2014] and the graph structure output by Minia was further simplified using Bifrost [Holley and Melsted, 2020] (Supplementary Section 2.5.2). As a result, we obtained a set of 747 unitigs with a total length of 373894 bp, an N50 of 840 bp, and a median length of 254 bp. We aligned the unitigs to the reference genome using BLAT [Kent, 2002] and found the alignment of one of the ten longest unitigs (of length 4887 bp) to span the expected breakpoint (Supplementary Figure 3, Supplementary Section 2.5.3). Within this unitig, a continuous subsequence of 766 bp that did not align to the reference genome was confirmed to be the sought variant (Supplementary Section 2.5.4).

## Discussion

In this paper, we introduce the efficient mapping tool MoleMap for linked- and long-read data and demonstrate its performance on simulated and real data sets in comparison to popular alignment and mapping tools.

MoleMap outputs a mapping score for each reported mapping that may be used in various ways. We may use the score as a quality weight factor in downstream analyses, or we may define a score threshold and limit the output to high-confidence mappings (Supplementary Section 3.6). We do not recommend immediately discarding all mappings with low scores, since there is some trade-off between precision and recall. In particular, we discourage using a score threshold when downstream analyses, such as alignment, ensure high precision.

We propose and demonstrate two main use cases for MoleMap. The first use case is a targeted analysis. Here, MoleMap serves as a pre-filter to detect reads from regions of interest and is followed by a standard alignment-based workflow restricted to the filtered reads. Assuming the speed-up factor of 8 observed in comparison to full read alignment on PacBio HiFi data in our evaluations (approx. 12 on ONT and linked reads), filtering with MoleMap requires 12.5% (8.33%) of the standard workflow’s running time. Consequently, we calculate that we may reduce the running time of a targeted analysis by using MoleMap if up to approximately 87.5% (91.66%) of the genome are of interest. For example, if all coding regions are of interest (less than 2% of the genome), we achieve a 6.9-fold speed-up (approx. 10-fold speed-up). Note that this is a conservative estimation, not accounting for superlinear downstream time savings, e.g., when sorting the output. In our second use case, MoleMap and other mapping tools can fully replace alignment with straightforward time savings. This applies to all downstream analyses that depend on mapping positions, such as local assembly (Section 3.6), but do not require a sequence assignment at the level of individual bases.

In all of our assessments, MoleMap performed significantly better outside the CenSat regions. The addition of highly repetitive regions in the recent T2T reference poses a challenge to MoleMap that we aim to address in the future. While CenSat and other less accessible regions of the genome lack extensive annotations, MoleMap is well suited to speed up data analysis on commodity hardware in diagnostic pipelines.

## Author contributions statement

B.K. conceived the project. R.L. and B.K. developed the molecule mapping approach. R.L. implemented MoleMap. M.S. provided patient data sets. R.L. and B.K. designed the experiments. T.K. locally assembled the non-reference sequence variant. R.L. conducted all other analyses. R.L. and B.K. analysed the results and wrote the manuscript. All authors approved the final manuscript.

## Supplementary Data statement

Supplementary Data are available at *Bioinformatics* online.

## Acknowledgments

We thank Brian Caffrey for his debugging assistance during the early development phase. Computation has been performed on the HPC for Research cluster of the Berlin Institute of Health.

## Data availability

The source code of MoleMap is available from GitHub, https://github.com/kehrlab/MoleMap/. The scripts used for testing and benchmarking MoleMap are available from GitHub, https://github.com/kehrlab/MoleMapScripts. The 10x chromium linked-read data of sample NA12878 was downloaded from AWS Amazon, http://s3-us-west-2.amazonaws.com/10x.files/samples/genome/2.1.4/NA12878\_WGS_v2/NA12878_WGS_v2_fastqs.tar. The ONT ultra-long read data of sample NA12878 by the Platinum Pedigree Consortium is available from AWS Amazon through GitHub, https://github.com/Platinum-Pedigree-Consortium/Platinum-Pedigree-Datasets. We do not have parents’ consent for publication of in-house patient sequencing data. The study has been approved by the Charité Ethics Committee (EA2/202/18).

## Footnotes

^1^https://support.10xgenomics.com/genome-exome/software/pipelines/latest/what-is-long-ranger

