## Supplementary for "MoleMap: fast alignment-free molecule mapping for long-read and linked-read sequencing data"

### 1 Supplementary methods

#### 1.1 Look-up cycle in an open-addressing index

An open-addressing index stores the keys or key identifiers of all indexed elements along with the index values. In our case, the keys are hash values of our minimizers and we store the first 16 bases of the indexed ($k$-mer) as identifiers. When querying a minimizer ($k$-mer) in our index using our hash function, the stored identifier is first analyzed. If the identifier is invalid, the minimizer is not in the index. If the identifier matches the queried minimizer, the reference positions of the minimizer can be retrieved from the index. If the identifier does not match the queried minimizer, the index bucket is occupied by another minimizer and the queried minimizer is stored elsewhere. This is called a collision. We use double hashing to resolve collisions, i.e., we re-hash the previous hash value by a bit-wise `XOR` between the previous index position and a bit-shifted version of the original minimizer. The identifier is then queried from the new hash position. This process is repeated until the correct index position is found.

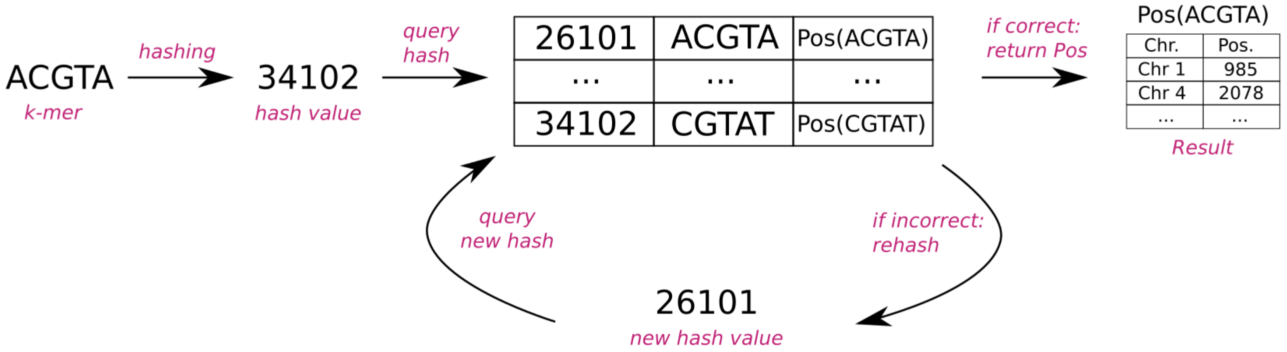

**Supplementary Figure 1:** Look-up cycle in an open-addressing k-mer index.

#### 1.2 Index table design in MoleMap

The index used by MoleMap consists of the identifier table, the directory table and the position table. To optimize the memory footprint, we set the sizes of the tables depending on the size of the indexed reference $l$, the $k$-mer length $k$, and the size of the minimizer windows $w$. We say that a minimizer is *active for $a$ bases* if it stays the minimizer in $a$ consecutive minimizer windows.

The identifier table stores an identifier of the contained minimizer and is used to resolve ambiguity from the open addressing. As the maximum amount of bases for which a minimizer can stay active is $w - k + 1$, on average a minimizer will be active for about half this long [1]. Hence, on average

$$S_I = \frac{l}{\frac{w-k+1}{2}} = \frac{2l}{w-k+1} \tag{1}$$

minimizers are necessary to cover the entire reference sequence and we choose this as the size of the identifier table. Because some minimizers will appear multiple times in the reference, fewer different minimizers than $S_I$ will have to be stored in the index. This circumstance ensures that placing minimizers in the identifier table results in only a small number of collisions.

The final number of minimizers per reference is hard to estimate, therefore we may adapt the index size after the initial calculation of minimizers. If necessary, we reset $S_I$ so that the identifier table is filled to at least 80%. This optimizes the memory requirements for indexing and mapping while still keeping the amount of collisions to a minimum.

The position table stores the reference positions of minimizers ordered by minimizers. The size of the position table is determined during index construction so that this table exactly fits the number of minimizers.

The directory table links the identifier table to the position table and is of size $S_I + 1$. If a minimizer is stored at index position $i$ of the identifier table, then position $i$ of the directory table points to the first entry in the position table that holds an occurrence of this minimizer. Position $i + 1$ of the directory table points to one entry past the last reference position of the minimizer (the first position of the next minimizer). The difference between position $i$ and $i + 1$ of the directory hence equals the number of occurrences of the minimizer in the reference genome. To save memory, we store no more than 16 bases of a minimizer as an identifier in the identifier table. For $k > 16$ this bears the theoretical risk of storing the positions of multiple minimizers in the same entry. However, as the whole base content is used for the hashing to determine the table position, collisions of this kind are extremely unlikely.

#### 1.3 Scaling factors $f(C)$

For long reads, we define the *scaling factor* $f(C)$ to account for the order of minimizer hits in the reference compared to the read: Given all pairs of minimizer hits $m_i$ and $m_{i+1}$ that are consecutive in the reference. We define $O$ as the difference between the number of consecutive minimizer hits in the reference where $o_m < o_{m+1}$ in the read, and the number of consecutive minimizer hits in the reference where $o_m \geq o_{m+1}$ in the read. As there are $|C| - 1$ pairs of consecutive minimizer hits, we set $f(C) = \frac{\text{abs}(O)}{|C|-1}$. If the order of minimizer occurrences in the read and reference is identical, $O$ will equal $|C| - 1$ and $f(C) = 1$; if the order is inconsistent, as expected for spurious minimizer hits, $O$ and $f(C)$ will be close to zero. In addition, the sign of $O$ indicates whether the read is mapped to the reference genome in forward or reverse orientation.

When mapping linked reads, MoleMap uses a scaling factor $f(C) = \frac{100}{L_C}$ that divides the sum in the cluster score by the length $L_C$ of the genomic region spanning the trimmed cluster. Dividing by $L_C$ prevents larger molecules from reaching higher mapping scores and is necessary because we do not have information about the number and length of mapped molecules. We multiply the score by $100$ to obtain a more tangible range of scores.

#### 1.4 Coverage analysis

For linked reads, MoleMap can use coverage profiles to dynamically adapt a local mapping score threshold (Supplementary Figure 2). More specifically, MoleMap estimates the average barcode coverage over the target genome. Then, after determining an upper bound for the minimum score threshold from the score histogram (Supplementary Section 3.5), MoleMap calculates a local score-specific barcode mapping coverage for fixed-size reference windows. Wherever the local barcode mapping coverage drops significantly below the expected uniform distribution, MoleMap locally adjusts the mapping score threshold to match the expected distribution.

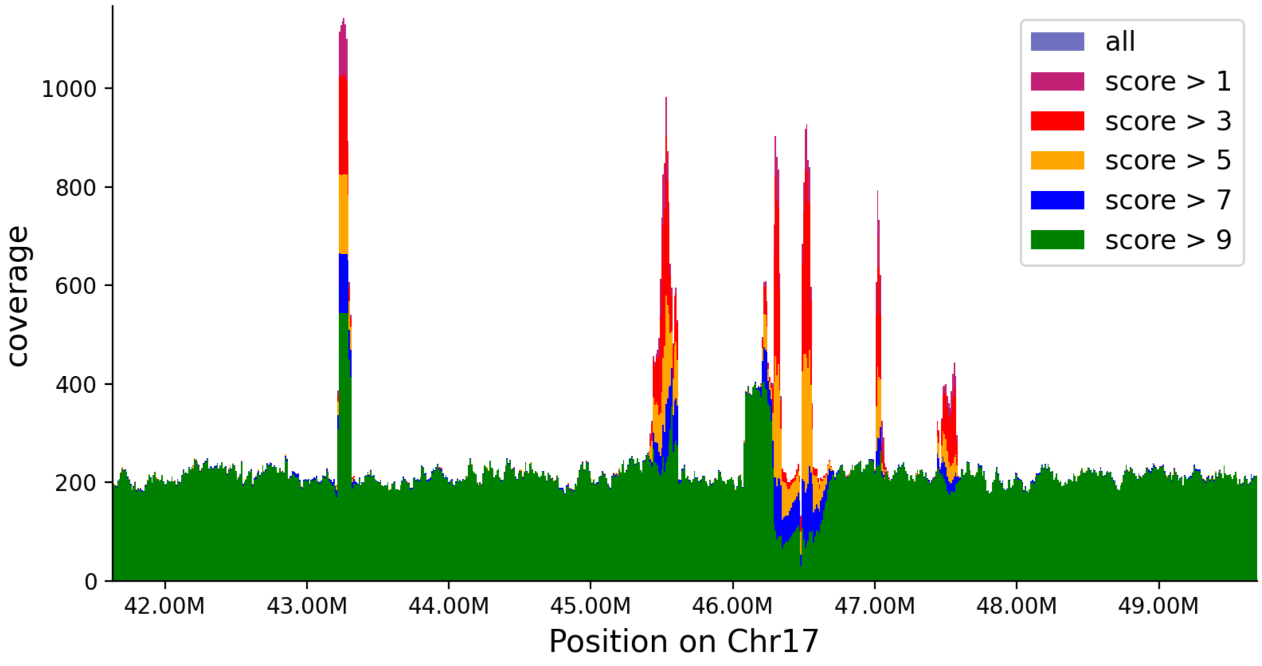

**Supplementary Figure 2:** Visualisation of a coverage profile.

This process allows for correctly distinguishing false positive mappings from low-score mappings caused by genomic events like segmental duplications. The fundamental assumption is that the coverage of correct mappings should be approximately uniformly distributed. Since short repeat elements, such as ALUs and LINEs, are not visible in these profiles, we assume that they have no significant impact on the mapping capabilities of MoleMap.

### 2 Data sets and command lines for evaluation

#### 2.1 Data sets for evaluation

##### 2.1.1 Real long-read data

**In-house PacBio HiFi data** We generated 14-fold coverage of PacBio HiFi data for a patient with congenital myopathy. Sequencing was performed by a commercial sequencing service provider on high-molecular weight DNA isolated from white blood cells using the classic salting out procedure [2]. Subsequent library preparation and sequencing was performed with an aimed for coverage of 30x on a PacBio Revio system.

**In-house Oxford Nanopore data.** We generated 50-fold coverage of ONT long-read data for a patient with both intellectual disability and muscular dystrophy carrying two previously identified diagnostic structural variants. Sequencing was performed by a commercial sequencing service provider on high molecular weight DNA isolated from white blood cells using the classic salting out procedure [2]. Subsequent library preparation and sequencing was performed with an aimed for coverage of 30x on an Oxford Nanopore PromethION flowcell.

**Public PacBio HiFi data** We downloaded PacBio HiFi reads published by the Platinum Pedigree Consortium [3] for sample NA12878. These data are already aligned and sorted. To obtain an unaligned, random subset of reads that corresponds to approximately 20-fold coverage, we randomized the order of the reads, selected the first 4 million reads after randomization, and converted the file to FASTQ format:

```
# Download of Platinum Pedigree HiFi data for NA12878
aws s3 cp --no-sign-request s3://platinum-pedigree-data/data/hifi/mapped/
    CHM13/NA12878.CHM13.haplotagged.bam NA12878.CHM13.haplotagged.bam

# Preparation of FASTQ files as input to MoleMap and aligners
samtools view -F 2048 NA12878.CHM13.haplotagged.bam | \
    awk 'BEGIN{srand();} {printf "%0.15f\t%s\n", rand(), $0;}' | \
    sort -n --parallel=4 --buffer-size=4G -T ./tmp/ | \
    head -n 4000000 | \
    awk '{print "@"$2; print $11; print "+"; print $12}' | \
    pigz -p 4 \
    > NA12878.CHM13.haplotagged.4M.fastq.gz
```

**Public Oxford Nanopore ultra-long data** We downloaded ONT ultra-long reads published by the Platinum Pedigree Consortium [3] for sample NA12878. These data are already aligned and sorted. To obtain an unaligned, random subset of reads that corresponds to approximately 20-fold coverage, we randomized the order of the reads, selected the first 1 million reads after randomization, and converted the file to FASTQ format:

```
# Download of Platinum Pedigree ONT UL data for NA12878
aws s3 cp --no-sign-request s3://platinum-pedigree-data/data/ont/mapped/
    CHM13/NA12878.minimap2.bam NA12878.minimap2.bam

# Preparation of FASTQ files as input to MoleMap and aligners
samtools view -F 2048 NA12878.minimap2.bam | \
    awk 'BEGIN{srand();} {printf "%0.15f\t%s\n", rand(), $0;}' | \
    sort -n --parallel=4 --buffer-size=4G -T ./tmp/ | \
    head -n 1000000 | \
    awk '{print "@"$2; print $11; print "+"; print $12}' | \
    pigz -p 4 \
    > NA12878.minimap2.1M.fastq.gz
```

##### 2.1.2 Real linked-read data

**In-house MGI stLFR data.** We generated 24-fold coverage of MGI stLFR data for the same patient with congenital myopathy as our in-house PacBio HiFi data. Sequencing was performed as a commercial sequencing service by the Beijing Genomics Institute (Hong Kong) on high molecular weight DNA isolated from white blood cells using the classic salting out procedure [2]. The stLFR libraray was prepared as published [4] and sequenced on a BGISEQ-500 machine.

**In-house TELL-Seq data.** We generated 46-fold coverage of TELL-Seq data for the same patient with congenital myopathy as our in-house PacBio HiFi and stLFR data. Sequencing was performed at the BIH Genomics facility on high-molecular weight DNA isolated from white blood cells using the classic salting out procedure [2]. The barcoded TELL-seq library [5] was prepared according to the manufacturer's (Universal Sequencing Technology, Carlsbad, CA, USA) specifications and sequenced on an Illumina NovaSeq 6000 system.

**10x Genomics.** We downloaded 10x linked read data published by 10x Genomics for sample NA12878/HG001. We selected the first sequencing lane corresponding to 36-fold coverage and corrected and trimmed barcodes using bcctools (`https://github.com/kehrlab/bcctools`).

```
# Download of NA12878 Germline Genome v2
wget http://s3-us-west-2.amazonaws.com/10x.files/samples/genome/2.1.4/
    NA12878_WGS_v2/NA12878_WGS_v2_fastqs.tar
tar xf NA12878_WGS_v2_fastqs.tar
rm NA12878_WGS_v2_fastqs.tar

# Preparation of FASTQ files as input to MoleMap and aligners
run_bcctools.sh -b bcctools -f fastq.gz -o 10x \
    NA12878_WGS_v2/NA12878_WGS_v2_S1_L001_R1_001.fastq.gz \
    NA12878_WGS_v2/NA12878_WGS_v2_S1_L001_R2_001.fastq.gz
```

##### 2.1.3 Simulated long-read data

We simulated long-read data with PBSIM from human chromosome 1 using the following commands. The reference file `chr1.fa` was extracted from T2T-CHM13v2.0.

**PacBio HiFi data:**

```
pbsim --strategy wgs --method qshmm \
    --depth 10 --genome chr1.fa \
    --qshmm pbsim3/data/QSHMM-RSII.model \
    --pass-num 10 --prefix HiFi_chr1
```

**ONT long-read data:**

```
pbsim --strategy wgs --method qshmm \
    --depth 10 --genome chr1.fa\
    --qshmm pbsim3/data/QSHMM-ONT.model \
    --prefix ONT_chr1
```

##### 2.1.4 Simulated linked-read data

We simulated linked-read data with LRSIM from human chromosome 21 with an without variation using the following commands. The reference file `chr21.fa` was extracted from T2T-CHM13v2.0.

**Without variation:**

```
simulateLinkedReads.pl \
    -r chr21.fa -o\
    -x 5 -t 90 -m 1\
    -0 0 -4 0 -7 0
```

**With variation:**

```
simulateLinkedReads.pl \
    -r chr21.fa -o\
    -x 5 -t 90 -m 1\
    -0 2 -4 10 -7 2
```

##### 2.1.5 Clustering of linked-read alignments into molecule mappings

LRSIM only reports the reference positions of simulated linked-reads but not the positions of the underlying molecules. To identify the positions of those simulated DNA molecules, we iterated the reads by position and defined a maximum cluster length threshold (300,000) and a maximum gap threshold (20,000) between read positions. We continuously added adjacent reads to the clusters until either threshold was exceeded. In this case we reported the cluster if it surpassed a minimum length threshold (10,000) and started the next cluster beginning with the next read. All parameters were chosen to match the default values of similar parameters used by MoleMap to achieve comparable results. The resulting intervals served as a truth set in the following analyses. The used python scripts are available at github (`https://github.com/kehrlab/MoleMapScripts`). To ensure a conservative evaluation of MoleMap, we use the same clustering procedure on the linked-read alignments output by the competing tools for evaluating their molecule mapping accuracy.

#### 2.2 Command lines

If not otherwise stated, tests were performed on an Intel Xeon E5-2650 v2 @2.60Ghz, 16 cores x2 threading. The **laptop analysis** was performed on a Lenovo T14s laptop with 8 cores (AMD Ryzen 7 PRO 5850U with 1901 MHz) and 16GB of main memory under WSL2 running Ubuntu 22.04.1 on Windows 10 Pro.

##### 2.2.1 Long-read mapping and alignment on real data

On real long-read data, we compared MoleMap to Minimap2, MashMap and BLEND. Furthermore, we ran Mapquick [6] but its reported mappings differed substantially from all other tools so that we decided to exclude it from evaluation. The index constructions listed in the following sections were not included in the running time and memory analyses.

**Input:**

```
T2T.fa      // Human reference genome
PacBio.fq   // PacBio HiFi sequencing data
ONT.fq      // ONT sequencing data
```

**MoleMap:**

```
molemap index T2T.fa --preset long -o T2T_Molemap_long

molemap mapLong PacBio.fq -t 32 -i T2T_Molemap_long -o PacBio_molemap.sam
molemap mapLong ONT.fq -t 32 -i T2T_Molemap_long -o ONT_molemap.sam
```

**Minimap2:**

```
minimap2 -t 32 -d T2T_HiFi.mmi -x map-hifi T2T.fa
minimap2 -t 32 -d T2T_ONT.mmi -x map-ont T2T.fa

minimap2 -t 32 -o PacBio_minimap2.paf T2T_HiFi.mmi PacBio.fq
minimap2 -t 32 -o ONT_minimap2.paf T2T_ONT.mmi ONT.fq
minimap2 -a -t 32 -o PacBio_minimap2.sam T2T_HiFi.mmi PacBio.fq
minimap2 -a -t 32 -o ONT_minimap2.sam T2T_ONT.mmi ONT.fq
```

**BLEND:**

```
blend -t 32 -d T2T_HiFi.bi -x map-hifi T2T.fa
blend -t 32 -d T2T_ONT.bi -x map-ont T2T.fa

blend -a -t 32 -o PacBio_blend.sam T2T_HiFi.bi PacBio.fq
blend -a -t 32 -o ONT_blend.sam T2T_ONT.bi ONT.fq
```

**MashMap:**

```
mashmap -r T2T.fa --saveIndex T2T_HiFi_mash -t 32 -q PacBio.fq \
    -o PacBio_mashmap.paf
mashmap -r T2T.fa --loadIndex T2T_HiFi_mash -t 32 -q PacBio.fq \
    -o PacBio_mashmap.paf

mashmap -r T2T.fa --saveIndex T2T_ONT_mash -t 32 -q ONT.fq \
    -o ONT_mashmap.paf
mashmap -r T2T.fa --loadIndex T2T_ONT_mash -t 32 -q ONT.fq \
    -o ONT_mashmap.paf
```

**Mapquick:**

```
mapquick -p pacBio_mapquick --threads 32 --reference T2T.fa PacBio.fq
mapquick -p ONT_mapquick --threads 32 --reference T2T.fa ONT.fq
```

##### 2.2.2 Long-read mapping and alignment on simulated data

We compared MoleMap to Minimap2, MashMap and BLEND on long read data simulated with PBSIM3 (Supplementary Section 2.1.3). We assume that the reference indexes have been constructed as specified in the previous section.

**Input:**

```
T2T.fa  // Human reference genome
PacBio_sim.fq     // Simulated PacBio HiFi sequencing data
ONT_sim.fq        // Simulated ONT sequencing data
```

**MoleMap:**

```
molemap mapLong PacBio_sim.fq -t 32 -i T2T_Molemap_long \
    -o PacBio_sim_molemap.sam
molemap mapLong ONT_sim.fq -t 32 -i T2T_Molemap_long \
    -o ONT_sim_molemap.sam
```

**Minimap2:**

```
minimap2 -a -t 32 -o PacBio_sim_minimap2.sam T2T_HiFi.mmi PacBio_sim.fq
minimap2 -a -t 32 -o ONT_sim_minimap2.sam T2T_ONT.mmi ONT_sim.fq
minimap2 -t 32 -o PacBio_sim_minimap2.paf T2T_HiFi.mmi PacBio_sim.fq
minimap2 -t 32 -o ONT_sim_minimap2.paf T2T_ONT.mmi ONT_sim.fq
```

**BLEND:**

```
blend -a -t 32 -o PacBio_sim_blend.sam T2T_HiFi.bi PacBio_sim.fq
blend -a -t 32 -o ONT_sim_blend.sam T2T_ONT.bi ONT_sim.fq
```

**MashMap:**

```
mashmap -r T2T.fa --saveIndex T2T_HiFi_mash -t 32 -q PacBio_sim.fq \
    -o PacBio_sim_mashmap.paf
mashmap -r T2T.fa --loadIndex T2T_HiFi_mash -t 32 -q PacBio_sim.fq \
    -o PacBio_sim_mashmap.paf

mashmap -r T2T.fa --saveIndex ONT_mash -t 32 -q ONT_sim.fq \
    -o ONT_sim_mashmap.paf
mashmap -r T2T.fa --loadIndex ONT_mash -t 32 -q ONT_sim.fq \
    -o ONT_sim_mashmap.paf
```

##### 2.2.3 Linked-read mapping and alignment on real data

On real linked-read data, we compared MoleMap to Long Ranger, BWA, EMA and Strobealign. We ran Strobealign in the default alignment mode and in "mapping only" mode. To run Long RAnger on stLFR and TELL-Seq data, we had to convert the sequencing data into a Long Ranger compatible format. The stLFR data was converted with the stLFR to Supernova pipeline (https://github.com/BGI-Qingdao/stlfr2supernova_pipeline). The TELL-SEq data was converted with the ust10x conversion tool from the General TELL-Seq Data Analysis Pipeline (https://universalsequencing.com/pages/tell-seq-software). We exclude EMA from the computaional performance analysis, as EMA uses BWA internally, so BWA's running time is a lower bound on EMA's running time.

##### Input:

```
T2T.fa                       // Human reference genome
10x.1.fq, 10x.2.fq           // barcode trimmed 10x Chromium sequencing data
stLFR.1.fq, stLFR.2.fq       // barcode trimmed stLFR sequencing data
tellSeq.1.fq, tellSeq.2.fq  // barcode trimmed tellSeq sequencing data

Special input data for Longranger:

10x_fq/10x_S1_L001_R1_001.fastq, \
10x_fq/10x_S1_L001_R2_001.fastq // raw 10x Chromium sequencing data

stLFR_fq/stLFR_S1_L001_R1_001.fastq, \
stLFR_fq/stLFR_S1_L001_R2_001.fastq //10x-like stLFR sequencing data

tellSeq_fq/tellSeq_S1_L001_R1_001.fastq, \
tellSeq_fq/tellSeq_S1_L001_R2_001.fastq // 10x-like tellSeq sequencing data
```

##### MoleMap:

```
molemap index T2T.fa --preset linked -o -o T2T_Molemap_linked

molemap maplinked 10x.1.fq 10x.2.fq -t 32 -i T2T_Molemap_linked \
    -o 10x_molemap.bed
molemap maplinked stLFR.1.fq stLFR.2.fq -t 32 -i T2T_Molemap_linked \
    -o stLFR_molemap.bed
molemap maplinked tellSeq.1.fq tellSeq.2.fq -t 32 -i T2T_Molemap_linked \
    -o tellSeq_molemap.bed
```

##### LongRanger:

```
mkdir -p T2Tref/fasta
ln -s T2T.fa T2Tref/fasta/genome.fa

longranger align --id 10x_out --fastqs 10x_fq --reference T2Tref \
    --localcores 32
mv 10x_out/outs/possorted_bam.bam 10x_longranger.bam

longranger align --id stLFR_out --fastqs stLFR_fq --reference T2Tref \
    --localcores 32
mv stLFR_out/outs/possorted_bam.bam stLFR_longranger.bam

longranger align --id tellSeq_out --fastqs tellSeq_fq --reference T2Tref \
    --localcores 32 // Longranger requires a modified whitelist within its\
    source folder to work on the converted tellSeq data.
mv tellSeq_out/outs/possorted_bam.bam tellSeq_longranger.bam
```

**BWA-MEM:**

```
bwa index T2T.fa

bwa mem -t 32 -o 10x_bwa.sam T2T.fa 10x.1.fq 10x.2.fq
bwa mem -t 32 -o stLFR_bwa.sam T2T.fa stLFR.1.fq stLFR.2.fq
bwa mem -t 32 -o tellSeq_bwa.sam T2T.fa tellSeq.1.fq tellSeq.2.fq
```

**EMA:**

```
ema align -1 10x.1.fq -2 10x.2.fq -t 32 -r T2T.fa -o 10x_ema.sam
```

**Strobealign:**

```
strobealign --create-index -t 32 T2T.fa 10x.1.fq 10x.2.fq
strobealign --create-index -t 32 T2T.fa stLFR.1.fq stLFR.2.fq
strobealign --create-index -t 32 T2T.fa tellSeq.1.fq tellSeq.2.fq

strobealign --use-index -o 10x_strobealign.sam -t 32 \
    T2T.fa 10x.1.fq 10x.2.fq
strobealign --use-index -o stLFR_strobealign.sam -t 32 \
    T2T.fa stLFR.1.fq stLFR.2.fq
strobealign --use-index -o tellSeq_strobealign.sam -t 32 \
    T2T.fa tellSeq.1.fq tellSeq.2.fq
```

**Strobealign in ”mapping only” mode:**

```
strobealign -x --use-index -o 10x_strobealign_x.paf -t 32 \
    T2T.fa 10x.1.fq 10x.2.fq
strobealign -x --use-index -o stLFR_strobealign_x.paf -t 32 \
    T2T.fa stLFR.1.fq stLFR.2.fq
strobealign -x --use-index -o tellSeq_strobealign_x.paf -t 32 \
    T2T.fa tellSeq.1.fq tellSeq.2.fq
```

##### 2.2.4 Linked-read mapping and alignment on simulated data

We tested Molemap to Long Ranger, BWA, EMA and Strobealign on 10x chromium data simulated with LRSIM (Supplementary Section 2.1.4). We assume that the reference indexes have been constructed as specified in the previous section.

**Input:**

```
T2T.fa  // Human reference genome
10x_sim.1.fq, 10x_sim.2.fq  \
// Simulated barcode trimmed 10x Chromium sequencing data

10x_sim_fq/10x_sim_S1_L001_R1_001.fastq, \
10x_sim_fq/10x_sim_S1_L001_R1_001.fastq \
// Simulated raw 10x Chromium sequencing data
```

**MoleMap:**

```
molemap maplinked 10x_sim.1.fq 10x_sim.2.fq -t 32 -i T2T_Molemap_linked \
    -o 10x_sim_molemap.bed
```

**LongRanger:**

```
longranger align --id 10x_sim_out --fastqs 10x_sim_fq --reference T2Tref \
    --localcores 32
mv 10x_sim_out/outs/possorted_bam.bam 10x_sim_longranger.bam
```

**BWA-MEM:**

```
bwa mem -t 32 -o 10x_sim_bwa.sam T2T.fa 10x_sim.1.fq 10x_sim.2.fq
```

**EMA:**

```
ema align -1 10x_sim.1.fq -2 10x_sim.2.fq -t 32 -r T2T.fa -o 10x_sim_ema.sam
```

**Strobealign:**

```
strobealign --use-index -o 10x_sim_strobealign.sam -t 32 \
    T2T.fa 10x_sim.1.fq 10x_sim.2.fq
```

**Strobealign in ”mapping only” mode:**

```
strobealign -x --use-index -o 10x_sim_strobealign_x.paf -t 32 \
    T2T.fa 10x_sim.1.fq 10x_sim.2.fq
```

##### 2.2.5 Exclusion of CenSat regions

We downloaded the CenSat annotation for the Human T2T genome from the UCSC table browser (https://genome.ucsc.edu/cgi-bin/hgTables). To exclude the CenSat regions from the alignments we used samtools with the following command:

```
Input:   CenSat.bed
         Alignment.sam
Command: samtools view --regions-file CenSat.bed \
             -U Alignment_noCenSat.sam Alignment.sam > /dev/null
```

#### 2.3 Linked-read mapping workflow

We compared the running time of full stLFR linked-read analysis workflows using MoleMap, Strobealign, BWA or Long Ranger. The workflows include data pre-processing, mapping/alignment and sorting. The output are mapped/aligned reads, from which regions of interest can be extracted efficiently. Reference index files were constructed according to Supplementary Section 2.2.3.

##### Input:

```
T2T.fa                            // Human reference genome
stLFR_raw.1.fq, stLFR_raw.2.fq   // stLFR sequencing data
```

**MoleMap:** The MoleMap workflow consists of barcode trimming and correction with bcctools, sorting of the corrected FASTQ input file with unix `sort` and mapping with MoleMap. Sorting with unix `sort` is included in the run_bcctools.sh script. To show the impact of the sorting, we display it separately in Figure 2D of the main text. The running time for barcode correction was measured without sorting, the and running time for sorting was calculated as the difference in running time when run with and without sorting.

```
scripts/run_bcctools.sh -o stLFR_corrected_sorted -f fastq -b ./bcctools \
    stLFR_raw.1.fq stLFR_raw.2.fq

molemap maplinked stLFR_corrected_sorted.1.fastq \
    stLFR_corrected_sorted.2.fastq -t 32 -i T2T_Molemap_linked \
    -o stLFR_molemap.bed
```

**Strobealign and BWA:** The alignment workflows for Strobealign and BWA include barcode trimming and correction with bcctools, alignment with Strobealign or BWA and sorting of the output files with samtools.

```
scripts/run_bcctools.sh -o stLFR_corrected -f fastq -b ./bcctools \
    stLFR_raw.1.fq stLFR_raw.2.fq

strobealign --use-index -o stLFR_strobealign.sam -t 32 T2T.fa \
    stLFR_corrected.1.fastq stLFR_corrected.2.fastq

bwa mem -t 32 -o stLFR_bwa.sam T2T.fa stLFR_corrected.1.fastq \
    stLFR_corrected.2.fastq

samtools sort -@ 31 stLFR_strobealign.sam > stLFR_strobealign.sorted.bam

samtools sort -@ 31 stLFR_bwa.sam > stLFR_bwa.sorted.bam
```

**Long Ranger:** Alignment with Long Ranger requires an additional pre-processing step which converts stLFR linked reads into the 10x Chromium linked-read format. Preprocessing was performed with the stLFR to supernova pipeline. Long Ranger trims and corrects barcodes and aligns the reads in one step. We sorted aligned reads with samtools.

```
awk '{if(NR>2) print($0)}' profile > profile_end
echo r1=\"{stLFR_raw.1.fq}\" > profile
echo r2=\"{stLFR_raw.2.fq}\" >> profile
cat profile_end >> profile
step_0_split_barode.sh
step_1_filter.sh
step_2_fake_10X_data.sh
mkdir fake10x
mv read-R1_si-TTCACGCG_lane-001-chunk-001.fastq.gz \
    fake10x/stLFR_S1_L001_R1_001.fastq
mv read-R2_si-TTCACGCG_lane-001-chunk-001.fastq.gz \
    fake10x/stLFR_S1_L001_R2_001.fastq

longranger align --id stLFR_longranger --fastqs fake10x \
    --reference T2Tref --localcores 32
mv stLFR_longranger/outs/possorted_bam.bam stLFR_longranger.sam

samtools sort -@ 31 stLFR_longranger.sam > stLFR_longranger.sorted.bam
```

#### 2.4 Targeted variant calling on ONT patient data

##### 2.4.1 Gene lists

**Muscular dystrophy:**

```
ACTA1, ANO5, B3GALNT2, BAG3, BIN1, CAPN3, CAV3, CHKB, COL6A1, COL6A2,
COL6A3, CPT2, CRYAB, DAG1, DES, DMD, DNAJB6, DNM2, DYSF, EMD, FHL1, FKRP,
FKTN, FLNC, GAA, GMPPB, ISCU, KLHL9, LAMA2, LDB3, LMNA, LPIN1, MTM1, MYH7,
MYOT, PNPLA2, POMGNT1, POMK, POMT1, POMT2, PYGM, RYR1, SECISBP2, SGCA,
SGCB, SGCD, SGCE, SGCG, SGCZ, SIL1, SMCHD1, TCAP, TIA1, TNNT1, TNPO3,
TOR1AIP1, TPM2, TPM3, TRAPPC11, TRIM32, TTN
```

**Intellectual disability:**

```
ACTB, ADNP, AHDC1, ANKRD11, ARID1B, ARX, AUTS2, ATRX, CASK, CDK13, CHD2,
CHD7, CLCN4, CREBBP, CTNNB1, DDX3X, DYNC1H1, DYRK1A, EHMT1, EP300, ERCC6,
FOXP1, GRIN2B, HUWE1, ITPR1, KAT6A, KAT6B, KCNQ2, KMT2A, KMT2D, MECP2,
MED12, MED13L, NALCN, NF1, NSD1, POGZ, PPP2R5D, PURA, PTPN11, SATB2, SCN2A,
SCN8A, SETD5, SETBP1, SLC6A1, SMC1A, SRCAP, STXBP1, SYNGAP1, TCF4, TCF20
```

**Conversion to bed:** We downloaded the NCBI RefSeq T2T gene predictions from the UCSC table browser and saved it as Homo_sapiens.T2T.gtf.gz (`https://genome.ucsc.edu/cgi-bin/hgTables`). We then extracted the gene positions from the annotation and extended the gene regions by $1000bp$ in each direction using the following command.

```
Input:  Homo_sapiens.T2T.gtf.gz
        geneList_ID.txt
        geneList_MD.txt
Command:
while read in; do
    grep "\"$in\"" <( zcat Homo_sapiens.T2T.gtf.gz ) -m 1 | \
    awk 'BEGIN{OFS=""} {print("chr",$1, "\t", $4, "\t", $5, "\t",\
    substr($14,2,length($14)-3))}' | \
    awk'{if(int($2)<1000) print($1,0,int($3)+1000,$4)} \
        {if(int($2)>=1000) print($1,int($2)-1000,int($3)+1000,$4)}'; \
done < <(cat geneList_MD.txt geneList_ID.txt)
```

##### 2.4.2 Variant calling workflow

```
Input:    [T2T.mmi] Minimap2 index of the T2T genome,
          [MoleMap_Index_T2T] MoleMap index of the T2T genome,
          [ONT_sample.fastq] Patient sequencing data,
          [Genelist_ID.bed] List of genes relevant for ID,
          [Genelist_MD.bed] List of genes relevant for MD

Output:   [SV-calls.vcf] Structural variants from regions of interest

Commands: cat Genelist_ID.bed Genelist_MD.bed > Genelist.bed
          molemap mapLong ONT_sample.fastq -t 32 -r Genelist.bed \
              -o Local_ONT_mapped.sam -i MoleMap_Index_T2T
          samtools fastq Local_ONT_mapped.sam > Local_ONT.fastq
          minimap2 -t 32 -a -o Local_ONT_realigned.sam T2T.mmi \
              Local_ONT.fastq
          samtools sort -@ 31 --write-index \
              -o Local_ONT_realigned.sorted.bam Local_ONT_realigned.sam
          sniffles --input Local_ONT_realigned.sorted.bam \
              --vcf SV-calls.vcf --threads 32
```

##### 2.4.3 Identified variants

```
chr12 115966387 Sniffles2.DEL.41SB N <DEL> 60 PASS PRECISE;SVTYPE=DEL; \
    SVLEN=-148429;END=116114815;SUPPORT=20;COVERAGE=49,29,18,31,46; \
    STRAND=+;STDEV_LEN=0.000;STDEV_POS=0.000;VAF=0.769 GT:GQ:DR:DV \
    0/1:4:6:20

chrX 32083165 Sniffles2.DEL.76S16 N <DEL> 60 PASS PRECISE;SVTYPE=DEL; \
    SVLEN=-164976;END=32248140;SUPPORT=24;COVERAGE=32,12,23,18,37; \
    STRAND=+;STDEV_LEN=0.000;STDEV_POS=0.000;VAF=1.000 GT:GQ:DR:DV \
    1/1:60:0:24
```

#### 2.5 Local assembly of a 766 bp non-reference sequence variant

##### 2.5.1 Linked-read extraction

```
Task:    Construct k-mer index of GRCh38
Input:   [GRCh38.fq] Human reference genome
Output:  [Index_GRCh38] Folder containing the bcmap k-mer index
Command: ./molemap index GRCh38.fq -o Index_GRCh38
```

```
Task:    Map barcodes of NA12878
Input:   [NA12878_linked_reads_1.fq, NA12878_linked_reads_2.fq],
         [Index_GRCh38]
Output:  [NA12878_mapped.bed] Barcode index produced by MoleMap
         [NA12878_read_index] Read index for NA12878 read files
Command: ./molemap map NA12878_linked_reads_1.fq \
             NA12878_linked_reads_2.fq \
             -i Index_GRCh38 \
             -o NA12878_mapped.bed \
             -b NA12878_read_index
```

```
Task:    Extract Barcodes of interest from barcode index
Input:   [NA12878_mapped.bed]
Output:  [Barcodes_NA12878_50K_region.txt]
Command: awk '{if($1=="chr17" \
             && (int($2)<=17831079+25000 \
             && int($3)>=17831079-25000)) \
             print($0)}' NA12878_mapped.bed \
             | awk '{if(int($5)>8) print($4)}' \
             > Barcodes_NA12878_50K_region.txt
```

```
Task:    Extract reads from read index
Input:   [NA12878_linked_reads_1.fq, NA12878_linked_reads_2.fq],
         [Barcodes_NA12878_50K_region.txt], [NA12878_read_index]
Output:  [NA12878_50K_region.fq]
Command: ./bcmap get \
             NA12878_linked_reads_1.fq \
             NA12878_linked_reads_2.fq \
             Barcodes_NA12878_50K_region.txt \
             -b NA12878_read_index \
             -o NA12878_50K_region.fq
```

##### 2.5.2 Linked-read assembly

```
Task:    Assembly of paired-end linked-reads.
Input:   [NA12878_50K_region.fq]
         File of interleaved pairs of linked-reads
Output:  [assembly_k121.unitigs.fa]
         A set of unitigs from the assembler's final iteration
Command: gatb \
            --12 NA12878_50K_region.fa \
            --no-scaffolding \
            --nb-cores 8 \
            > logs/assembly.log 2>&1
```

```
Task:    Simplifying the set of unitigs in their de Bruijn Graph
         representation, i.e., removing tips and singletons.
Input:   [assembly_k121.unitigs.fa]
         The set of unitigs from the previous step.
Output:  [assembly_k121.unitigs.fa.gfa]
         A simplified de Bruijn Graph in GFA format.
Command: Bifrost build \
            -r assembly_k121.unitigs.fa \
            -t 8 \
            -k 121 \
            -m 81 \
            --clip-tips \
            --del-isolated \
            -o assembly_k121.unitigs.fa
```

```
Task:    Extract unitigs from simplified de Bruijn Graph.
Input:   [assembly_k121.unitigs.fa.gfa]
         The simplified de Bruijn Graph from the previous step.
Output:  [assembly_k121.unitigs.bifrost.fa]
         The unitigs of the simplified de Bruijn Graph in FASTA format.
Command: awk '$0 ~ /^S/ {print ">"$2"\n"$3}' \
            assembly_k121.unitigs.fa.gfa \
            > assembly_k121.unitigs.bifrost.fa
```

##### 2.5.3 Assembly validation

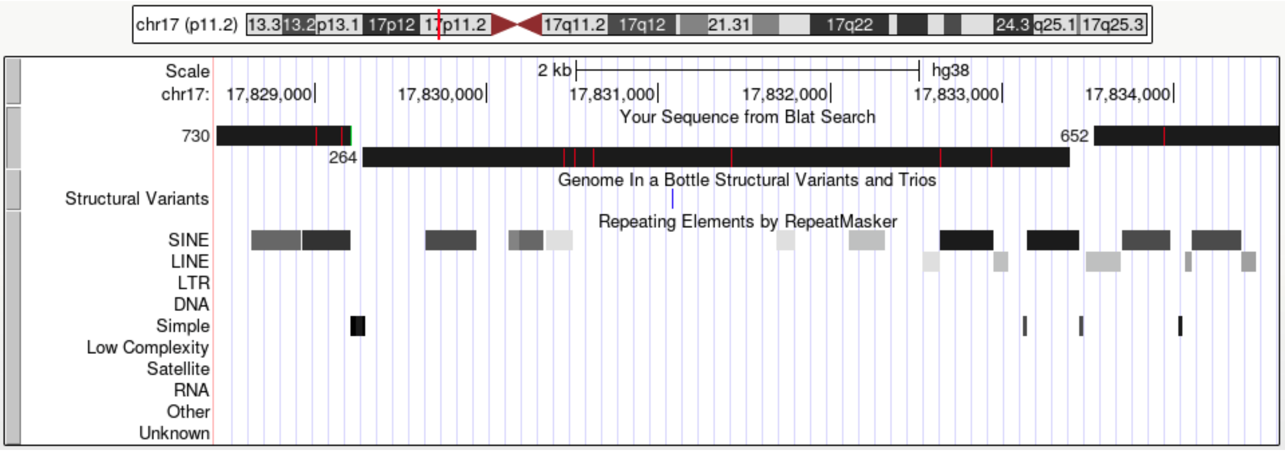

**Supplementary Figure 3:** Alignment of the unitigs from the local assembly to the HG38 reference using the UCSC web application of BLAT [7]. The unitig with the ID 264 spans the target location at chr17:17,831,079. The blue mark in the *Structural Variants* track denotes the sought variant as previously reported in the Genome in a Bottle (GIAB) Structural Variants callset.

##### 2.5.4 Sequence validation of the non-reference sequence variant.

We performed a local sequence alignment of the continuous subsequence of unitig 264 that does not align to the reference genome to the insertion sequence at breakpoint chr17:17,831,079 reported in the GIAB callset. The two sequences, both of 766 bp length, align with 100% identity and similarity. The alignment was computed using the EMBOSS Water web application [8] of the Smith-Waterman algorithm for local sequence alignment. The web application's output is documented below.

```
########################################
# Program: water
# Rundate: Thu  6 Jan 2022 16:01:26
# Commandline: water
#    -auto
#    -stdout
#    -asequence emboss_water-I20220106-160436-0340-65113364-p2m.aupfile
#    -bsequence emboss_water-I20220106-160436-0340-65113364-p2m.bupfile
#    -datafile EDNAFULL
#    -gapopen 10.0
#    -gapextend 0.5
#    -aformat3 pair
#    -snucleotide1
#    -snucleotide2
# Align_format: pair
# Report_file: stdout
########################################

#=======================================
#
# Aligned_sequences: 2
# 1: chr17_17831079_pbsv.INS.40992
# 2: 264
# Matrix: EDNAFULL
# Gap_penalty: 10.0
# Extend_penalty: 0.5
#
# Length: 766
# Identity:     766/766 (100.0%)
# Similarity:   766/766 (100.0%)
# Gaps:           0/766 ( 0.0%)
# Score: 3830.0
#
#
#=======================================
```

### 3 Supplementary results

#### 3.1 Precision and Recall calculation

For computing recall and precision, we define a predicted molecule mapping as a true positive if the original interval in the truth set overlaps the predicted mapping with start and end positions each deviating by less than 20,000 bases. A predicted mapping that has no such equivalent in the truth set is defined as a false positive, and a true interval without a predicted equivalent is defined as a false negative. Furthermore, we require a one-to-one correspondence for true positives, meaning that a true interval can only qualify one predicted mapping as a true positive.

#### 3.2 Computational Performance

The following tables contain the exact running times and memory requirements that are plotted in Figure 2 of the main article.

**Supplementary Table 1:** Running times [min] and memory consumption [GB] on PacBio-HiFi data.

| # Threads | 1 | | 8 | | 16 | | 32 | |
|---|---|---|---|---|---|---|---|---|
| | time | mem | time | mem | time | mem | time | mem |
| MoleMap | 202 | 3.84 | 38 | 3.87 | 20 | 3.88 | 10 | 3.96 |
| Minimap2 align | 1404 | 13.53 | 293 | 18.53 | 195 | 21.31 | 192 | 24.88 |
| Minimap2 map | 437 | 11.87 | 69 | 12.26 | 38 | 12.45 | 32 | 12.58 |
| BWA | - | - | - | - | - | - | 2541 | 8.67 |
| Blend | 1349 | 8.13 | 209 | 14.52 | 119 | 19.06 | 106 | 26.92 |
| MashMap | - | - | 377 | 40.52 | 218 | 40.52 | 136 | 40.53 |

**Supplementary Table 2:** Running times [min] and memory consumption [GB] on ONT data

| # Threads | 1 | | 8 | | 16 | | 32 | |
|---|---|---|---|---|---|---|---|---|
| | time | mem | time | mem | time | mem | time | mem |
| MoleMap | 597 | 6.5 | 122 | 6.56 | 53 | 6.62 | 29 | 6.76 |
| Minimap2 align | 6572 | 12.4 | 1460 | 16.4 | 619 | 19.5 | 390 | 22.4 |
| Minimap2 map | - | - | 868 | 10.34 | 442 | 10.99 | 233 | 12.22 |
| Blend | - | - | 1694 | 19.77 | 875 | 23.6 | 366 | 26.9 |
| MashMap | 3030 | 34.21 | 649 | 34.5 | 259 | 34.8 | 226 | 35.45 |

**Supplementary Table 3:** Running times [min] and memory consumption [GB] on stLFR data

| # Threads | 1 | | 8 | | 16 | | 32 | |
|---|---|---|---|---|---|---|---|---|
| | time | mem | time | mem | time | mem | time | mem |
| MoleMap | 106 | 2.72 | 17 | 2.81 | 12 | 2.83 | 9 | 2.85 |
| BWA | 8349 | 6.01 | 1854 | 7.81 | 923 | 9.31 | 487 | 12.9 |
| Longranger | - | - | - | - | - | - | 1493 | 57.1 |
| Strobealign align | 2397 | 15.03 | 452 | 15.38 | 206 | 15.82 | 100 | 16.71 |
| Strobealign map | 472 | 15.03 | 164 | 15.16 | 103 | 15.39 | 47 | 15.89 |

**Supplementary Table 4:** Mapping Workflow running times on stLFR data [min]

| | Format Conversion | Barcode Correction | Mapping/Alignment | Sorting |
|---|---|---|---|---|
| MoleMap | - | 172 | 17 | 107 |
| Strobealign | - | 172 | 359 | 126 |
| BWA | - | 172 | 774 | 126 |
| Longranger | 744 | - | 1842 | 126 |

#### 3.3 Mapping quality with introduced variation.

We tested MoleMap on simulated linked-read data with and without introducing variation (supplementary section 2.1.4) to test the robustness of the mappings to small changes. Additional variation had no distinct impact on mapping quality.

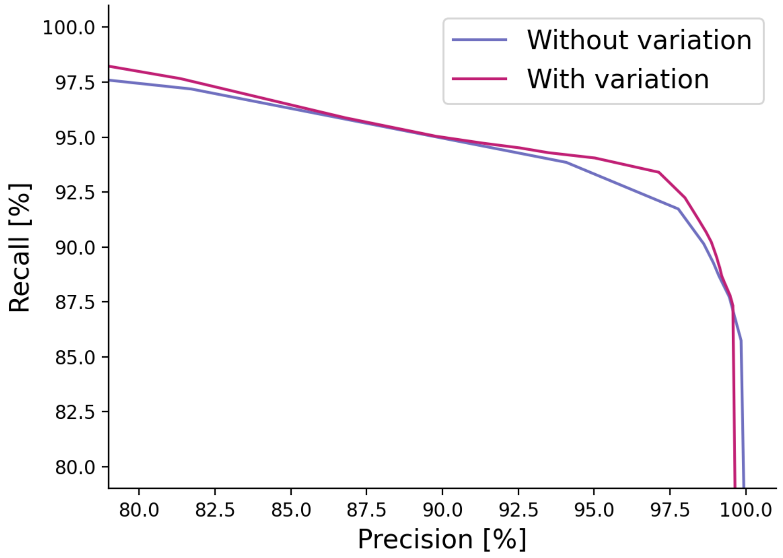

**Supplementary Figure 4:** Precision-recall curve for MoleMap with and without introduced variation in the simulated data.

#### 3.4 Impact of minimizers on computational performance

The mapping approach in MoleMap owes its computational performance to the use of minimizers. We quantified the impact of minimizers in MoleMap on the construction time and size of the $k$-mer index build for the reference genome as well as on molecule mapping.

We constructed the $k$-mer index of the T2T reference using 31-mers and minimizer window sizes $w$ spanning from 41 to 61 as well as 17-mers with $w$ from 21 to 25. The time for constructing and the memory required by the index (Supplementary Table 5) scales according to the number of minimizers that are required to cover the reference (Supplementary equation (1)).

For the same parameters, we mapped the stLFR and HiFi data sets (Supplementary Section 2.1.2, 2.1.1) using the T2T indexes. The impact on the memory consumption of the mapping is is dominated by the size of the index. The impact on the running time of mapping is even squared compared to indexing since both the number of evaluated minimizers and the number of hits per evaluated minimizer scale up.

**Supplementary Table 5:** Impact of minimizer window size $w$ on wall clock time [min] and memory usage [GB] of MoleMap on real pacBio HiFi and stLFR data.

| $k = 17$ | $w = 21$ | | $w = 23$ | | $w = 25$ | |
|---|---|---|---|---|---|---|
| | time | mem | time | mem | time | mem |
| molemap index | 52:56 | 15.4 | 33:39 | 11.29 | 29:55 | 8.94 |
| molemap mapLong | 16:42 | 10.59 | 10:13 | 8.01 | 8:17 | 6.45 |
| $k = 31$ | $w = 41$ | | $w = 51$ | | $w = 61$ | |
| | time | mem | time | mem | time | mem |
| molemap index | 29:50 | 7.58 | 12:15 | 4.29 | 9:40 | 3.11 |
| molemap mapLinked | 16:11 | 24.68 | 10:01 | 21.77 | 8:23 | 20.59 |

#### 3.5 Impact of maximum gap threshold

We measured the impact of the maximum-gap-size parameter on the mapping accuracy using simulated read data (Supplementary section 2.1.3 and 2.1.4). Our tests show that the mapping quality is generally sensitive to this parameter (Supplementary Figure 5) but stable over a large range. A default values of 2,000 for long-reads and 20,000 for linked-reads were picked, based on this assessment. We note that different data sets may have different optimal values.

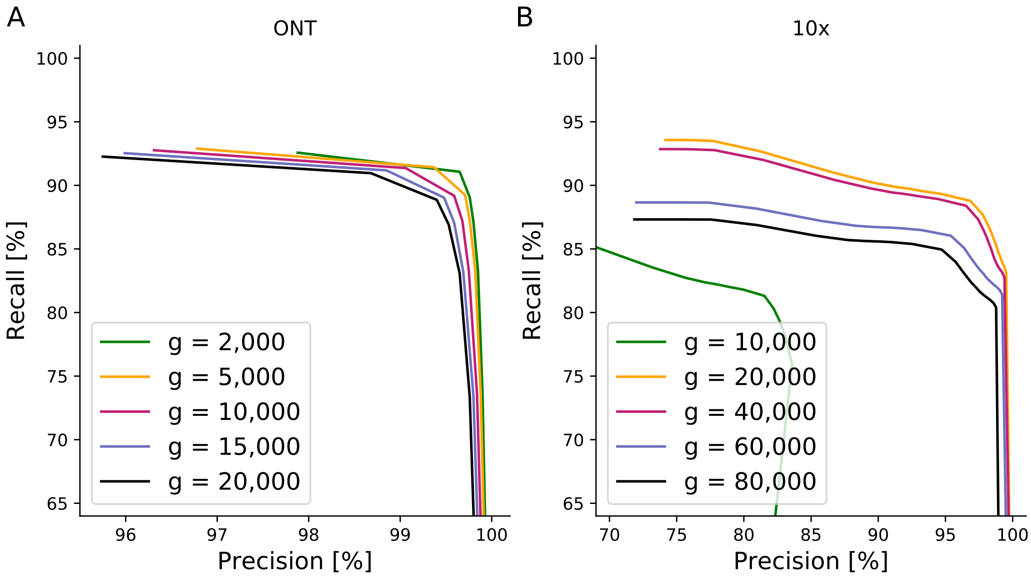

**Supplementary Figure 5:** Impact of the maximum gap size parameter $g$ on precision and recall on simulated ONT long-reads and 10x linked-reads.

##### 3.6 MoleMap's score separates correct from spurious mappings

On the simulated data, we observed that higher mapping score thresholds lead to a steadily increasing precision (Figure 3). This observation indicates that our mapping score can indeed distinguish true positive from false positive barcode mappings. However, the range of scores is expected to vary between different data sets and different reference genomes. For example, a larger reference with fewer unique or rare $k$-mers will lead to lower scores overall (Methods). A score threshold that works well for one data set or reference genome may not be appropriate for a different setup.

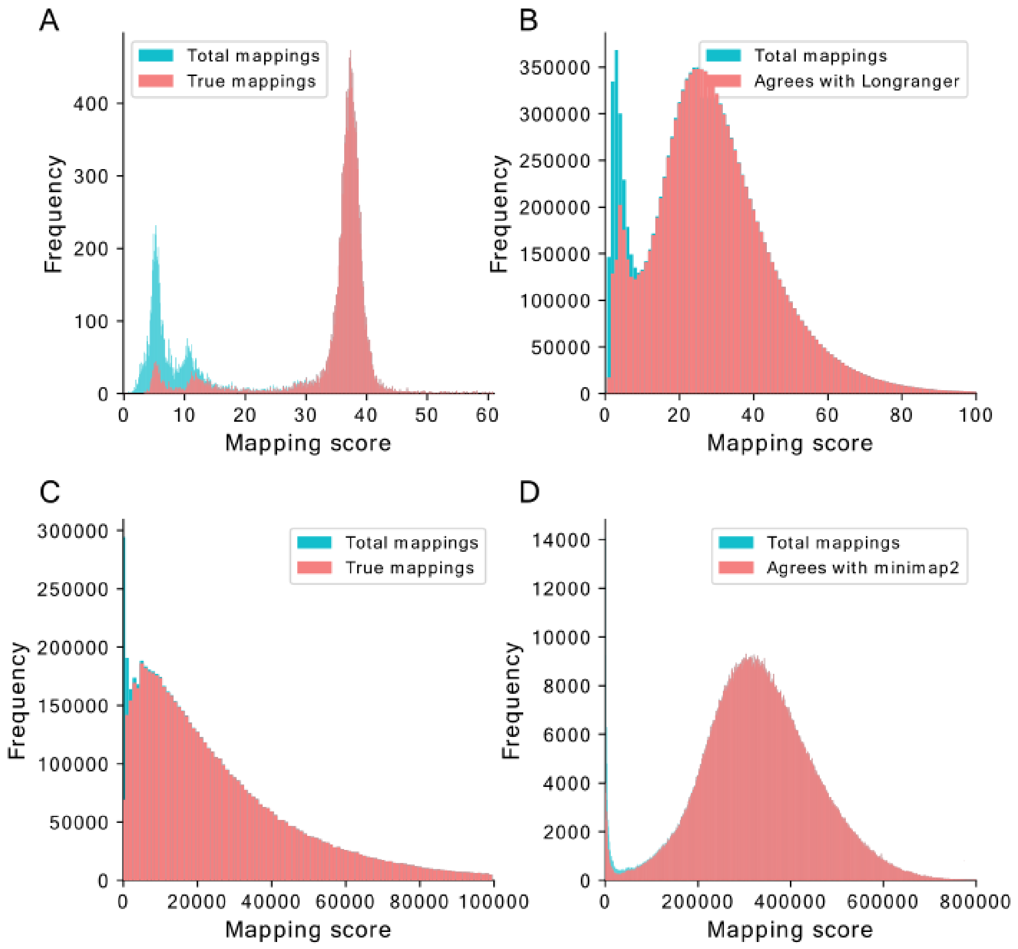

**Supplementary Figure 6:** Mapping score distribution for simulated linked read data (A), real 10x genomics linked reads (B), simulated PacBio HiFi long reads (C) and real PacBio HiFi long reads (D).

To provide guidance on reasonable score thresholds, we examine the score distributions for both simulated and real data sets (Supplementary Figure 6). Thorough analysis suggests that the score distribution follows a common pattern: one peak of unreliable, low scores and one peak of reliable, higher scores with a dip in-between. Simulated data (Supplementary Figure 6A,C) confirm a low true positive rate for the first peak and a high true positive rate for the second peak. On real data (Supplementary Figure 6B,D), the agreement with LongRanger and Minimap2 suggests the same separation. By determining the minimum between the two peaks, we can infer a score threshold for reliable barcode mappings on real data, e.g., a minimum mapping score of 10 in Supplementary Figure 6B and a minimum mapping score of 20,000 in Supplementary Figure 6D.
